# Erosion of regenerative regulation: age-associated shifts in the skeletal muscle fiber epigenome and transcriptome

**DOI:** 10.64898/2026.08.19.744884

**Authors:** Keagan G. Moo, Peter Orchard, Arushi Varshney, Ricardo D’Oliveira Albanus, Nandini Manickam, Leena Kinnunen, Timo A. Lakka, Jouko Saramies, Markku Laakso, Jaakko Tuomilehto, Karen L. Mohlke, Michael Boehnke, Laura Scott, Heikki A. Koistinen, Francis S. Collins, Stephen C. J. Parker

## Abstract

Skeletal muscle aging is characterized by the deterioration of muscle function, which can lead to negative quality-of-life outcomes including frailty and sarcopenia. While understanding the mechanisms of this process is increasingly important as the global population ages, previous molecular studies of skeletal muscle aging have been limited by statistical power and cell type resolution. In this study, we analyzed single-nucleus gene expression and chromatin accessibility data from 287 human skeletal muscle samples from individuals aged 20-79 years to explore sex- and cell type-specific aging effects. Across 467,126 nuclei from 13 cell types, we identify 384 age-associated genes and 4,061 age-associated chromatin regions. These age-associated molecular features are enriched for functional pathways, including metabolic processes, cell-to-cell communication, and senescence Kyoto Encyclopedia of Genes and Genomes KEGG terms. Age-associated closing chromatin was more common across fiber types and sexes than opening chromatin, and was enriched in active enhancer regions while depleted for active transcription start sites. We observe enrichment for specific transcription factor motifs in closing chromatin, including those of glucocorticoid and androgen receptors, both of which play a key role in the maintenance of healthy skeletal muscle. Together, these findings identify an age-associated regulatory shift, largely invisible in matched transcriptomic data, characterized by closing chromatin which reduces accessibility to hormone receptor binding sites and enhancer regions in the muscle fiber epigenome.

## Introduction

Skeletal muscle is the largest organ in the human body by mass, accounting for about 40%^1^ of body weight, consisting predominantly of multi-nucleated muscle fibers. These muscle fibers include ‘slow-twitch, oxidative type 1 muscle fibers and ‘fast-twitch’ glycolytic type 2a and 2x muscle fibers. Muscle mass peaks around age 30, after which a functional decline^2,3^, driven primarily by type 2 fiber loss^4^, increases the risk of many health complications including frailty^5^, falls^6^, insulin resistance^7^, immune dysregulation^8^, and death^9,10^. The molecular mechanisms driving this decline remain incompletely understood.

Prior studies examining age effects on the transcriptomic and epigenomic landscape of skeletal muscle either profile bulk tissue^11–14^, which obscures cell type-specific differences, or else have small sample sizes (<35 participants)^13,15,16^, limiting robust identification of age-associated molecular features.

Furthermore, most studies only include individuals from one sex or lack sex-stratified analyses. This represents an opportunity to build on existing evidence of sex-based differences in cell-type composition^17^, gene expression^17^, and functionality^1,18^ in human skeletal muscle to explore age effects at a single nucleus level.

In this study we leverage single-nucleus RNA and ATAC sequencing (snRNA-seq and snATAC-seq) from up to 287 human *vastus lateralis* samples^19^ to identify age-associated compositional, transcriptomic, and epigenomic shifts in different skeletal muscle cell types. Based on these age-associated changes in molecular features, we then test for enrichment of KEGG pathway genes, chromatin states, and transcription factor motifs, to give us a greater understanding of the mechanisms which underlie the process of aging in skeletal muscle tissue.

## Results

### Skeletal muscle nuclei type proportions change with age

To characterize aging-related changes in human skeletal muscle we used snRNA-seq and snATAC-seq on frozen *vastus lateralis* muscle biopsies from 287 donors, 123 female and 164 male, between 20 and 79 years of age (Fig 1A, B)^19^. Clustering identified 13 cell types. The majority of nuclei were from muscle fibers (33.8% type 1, 19.7% type 2a, and 15.4% type 2x) (Fig. 1C). Other cell types included, in order of abundance, endothelial, fibro-adipogenic progenitor (FAP), smooth muscle, T-cell, neuronal, neuromuscular junction, mixed-muscle fiber, satellite cell, adipocyte, and macrophage nuclei^19^.

**Figure 1:**
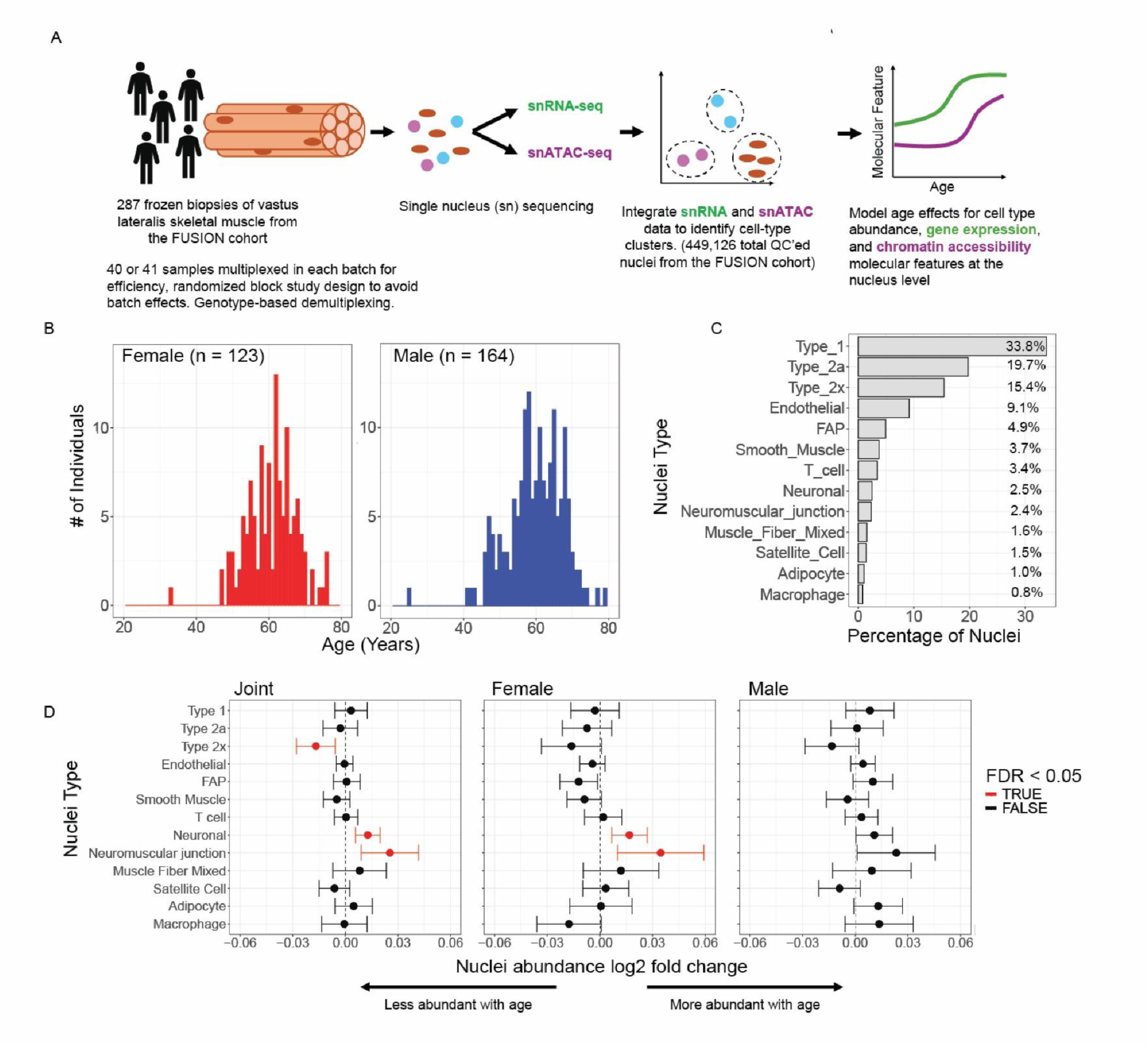
Age-associated differences in nuclei-type abundance in human skeletal muscle. a.) Overview of study design. Figure is based on Varshney, A et al. bioxrv (2024) now accepted at Nature Genetics b.) Sex specific donor age distribution and count. c.) Skeletal muscle nuclei-type abundance. d.) Association between each nuclei type’s abundance and age, using combined RNA and ATAC nuclei counts. Error bars represent 95% confidence intervals. Cell types with an FDR < 0.05 significant association between abundance and age are colored red. Joint = combined analysis of male and female samples.

To quantify age-associated changes in skeletal muscle composition, we tested for association between age and cell type abundance^20^. We identified a significant (FDR < 0.05) decrease of type 2x fiber nuclei alongside a significant increase in neuronal and neuromuscular junction nuclei (Fig 1D). To determine if aging affects skeletal muscle composition in a sex-specific manner, we repeated the analysis on female and male samples separately. In the female analysis both the neuronal and neuromuscular junction increases in nuclei abundance remained significant, but no other significant changes were identified in the sex specific analyses (Fig 1D).

### Broad age-associated changes to the muscle fiber transcriptome

To understand how the human skeletal muscle transcriptome changes with age, we tested for associations between gene expression and age using either all samples (“joint analysis”) or stratifying by sex. We used a linear mixed model (LMM) and adjusted for both biological and technical covariates (Methods). A pseudobulk approach using DESeq2 returned highly similar results (Fig S2B). The joint analysis identified 216, 384, and 344 significantly (FDR < 0.05) age-associated genes in type 1, type 2a, and type 2x fibers respectively. The sex-stratified analyses identified 184, 163, and 81 age-associated genes in females and 37, 152, and 147 age-associated genes in males (4, 20, and 13 genes FDR < 0.05 in both sex-specific analyses respectively) (Fig 2A). The less abundant cell types yielded < 50 total age-associated genes (Supplementary Table 1), therefore we focus on the three muscle fiber types for the remainder of this study (Fig S1A). Upregulated and downregulated age-associated genes were fairly evenly spread across the fiber types and sexes (Fig 2A). While genes meeting the significance threshold for age-association differed by sex and fiber type, the direction of age association for significant genes was broadly concordant across sex (72% - 81%) and fiber type (86% - 96%) (Fig 2B, Fig S2A). Collectively, these results suggest that while age-associated transcriptome changes occur in sex and fiber type-specific ways, the broad directional shifts in gene expression are similar across sexes and fiber types.

**Figure 2:**
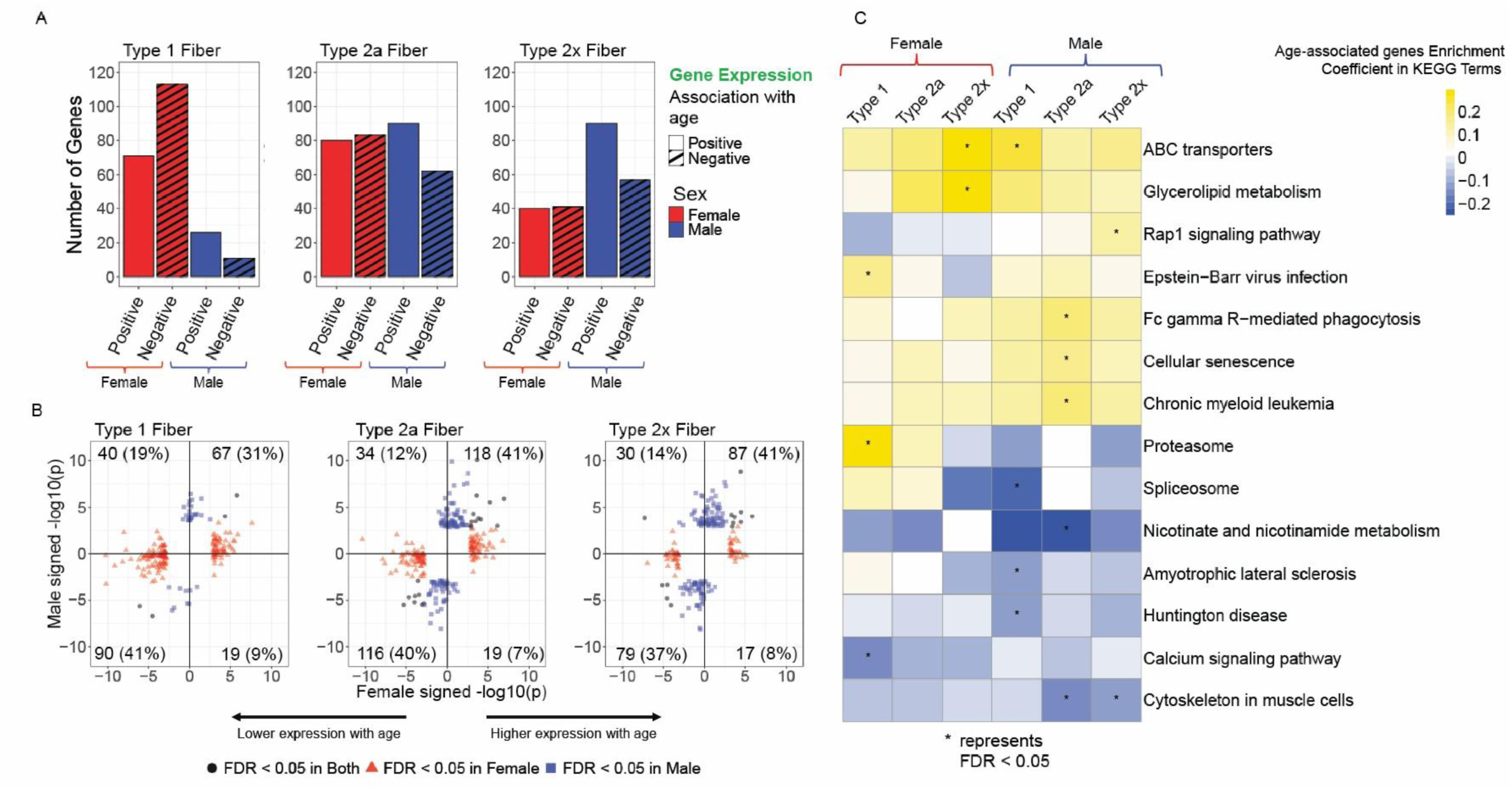
Age-associated changes in gene expression and pathway enrichment in human skeletal muscle fiber. a.) Number of genes identified as significantly age associated, by fiber type, sex, and direction of association with age. b.) Age-associated expression in males vs age-associated expression in females, for genes with age-associated expression in at least one sex. Positive x- and y-axis values represent higher expression with age, while negative values represent lower expression with age. c.) Heatmap of tested gene sets with FDR < 0.05 enrichment or depletion in at least one set (Sex and fiber type combination) of age-associated genes.

We compared our findings to those of Kedlian et al.^16^, who used an LMM with a local true sign rate (LTSR) significance threshold of > 0.9 to analyze differentially expressed genes in 17 intercostal muscle samples from young (20-40 years) versus old (60-75 years) subjects. Though Kedlian et al did not report nominal p-values for all genes tested, we can use the LTSR values assigned to genes they identified as significantly age-associated to compare to our nominal p-values. Of the genes Kedlian et al identified, 58-64% showed the same direction of effect in our data (regardless of significance), though few (16% or 110 genes) were significantly age-associated in our analysis^16^ (Fig S3A).

### Muscle age-associated genes are enriched for pathways involving muscle stress and immune signaling

Next we tested KEGG gene sets for enrichment of age-associated genes to identify biological pathways being associated with the aging process^21–24^. Most significantly (FDR < 0.05) enriched or depleted terms showed a consistent direction of effect between male and female fibers including cellular senescence, ABC transporters, and glycerolipid metabolism (Fig 2C) (Supplementary Table 2). In contrast, the proteasome term, our most significantly enriched term, was enriched for age-associated genes in female type 1 fiber while trending towards depletion for age-associated genes in male fiber types (Fig 2C). This enrichment for age-associated genes in the proteasome gene set was driven by significant increases in PSME4 and PSMD11 subunit expression in female type 1 fiber. Taken together, these results reiterate the similarities in age-associated transcriptomic changes across sex and fiber type and suggest that age-associated changes emphasize metabolic pathways, cell senescence, and cell-to-cell signalling.

### Age-associated remodeling of the muscle fiber epigenome

To explore how aging impacts the chromatin landscape in skeletal muscle, we used snATAC-seq data to test for association between chromatin accessibility and age. Starting with 149,751 - 183,850 previously identified open chromatin regions (OCRs) per fiber type^19^, we tested 36,075 - 44,469 OCRs with at least 1 count in 10% of a given nuclei type for age association. We used generalized linear mixed models and adjusted for both technical and biological covariates (Methods). A pseudobulk approach using DESeq2 returned highly similar results (Fig S5B). The joint analysis identified 831, 4,061, and 1,198 significantly (FDR < 0.05) age-associated OCRs in type 1, type 2a, and type 2x muscle fiber respectively. The sex-stratified analyses identified 319, 753, and 23 age-associated OCRs in females and 47, 1,356, and 365 age-associated OCRs in males (1, 141, and 1 OCRs FDR < 0.05 in both sex-specific analyses respectively) (Fig S4A) (Supplementary Table 3). We observed that for a given fiber type and sex, most (68% - 72%) age-associated OCRs showed a negative association between accessibility and age (i.e. were closing with age) (Fig 3A). We identified many more age-associated OCRs in type 1 fibers in females (n=319) compared to males (n=47), but in type 2x fibers more were identified in males (n=365) than females (n=23). Similar to our age-associated gene expression results, while few age-associated OCRs were significant in both sexes, the direction of regulation was broadly consistent (80% - 91%), with the type 1 fiber comparison showing the lowest concordance (Fig 3B). Within each sex, age-associated OCRs showed a consistent direction of association with age across fiber types (Fig S5A). These results suggest broad remodeling of the muscle fiber epigenome, especially in type 2a fibers, with similar aging patterns across sex and fiber type.

**Figure 3:**
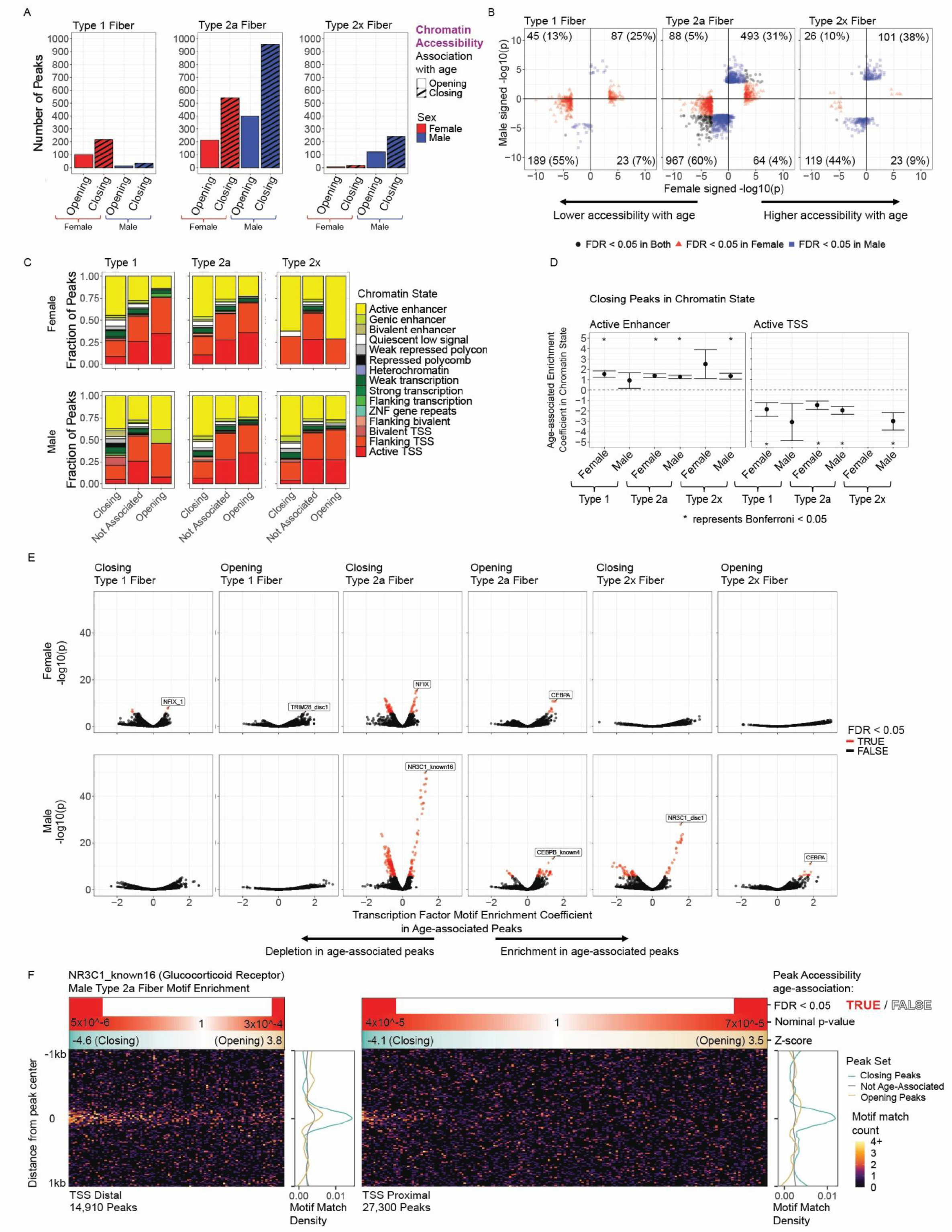
Age-associated changes in chromatin accessibility, state, and pathway enrichment in human skeletal muscle fiber. a.) Number of OCRs identified as significantly (FDR < 0.05) age associated counted by fiber type, sex, and association with age. b.) Signed -log10 p-values (+ more expression with age, - less expression with age) of differential chromatin accessibility in each fiber type between the female and male analyses. c.) Ratio of OCRs overlapping with Roadmap chromatin states. OCRs are either from significantly (FDR < 0.05) age-associated sets or from a non-age associated set with nominal p-value > 0.5 in all sex and fiber type combinations. d.) Enrichment coefficients and confidence intervals for active enhancer and active TSS in age-associated closing chromatin for each fiber type and sex combination. Due to the extremely low number of significantly age-associated peaks detected in female type 2x fiber and male type 1 fiber some logistic regression results produce extreme negative values as a result of encountering no overlapping OCRs (e.g. female type 2x fiber age-associated OCRs overlapping with the active TSS chromatin state). e.) Per sex and fiber type volcano plots of enrichment results for transcription factor motifs in age-associated OCRs. Transcription factor motifs with Bonferroni-adjusted p-value < 0.05 enrichment or depletion have been colored red. f.) Density of predicted TF binding sites for NR3C1 (glucocorticoid receptor) in OCRs (+/-1 kb around OCR center). OCRs are ordered along the heatmaps x-axis by their direction of association with age (z-score of the model coefficient), and separated into TSS distal and TSS proximal OCR subplots. Heatmaps represent predicted binding site density (black-purple = low density, yellow-white = high density). The teal-to-brown gradient plot represents OCR accessibility age-association z-score. The white and red plot indicates whether the plotted OCRs’ accessibility are significantly (FDR < 0.05) associated with age (red) or not (white). The white-to-red gradient plot gives a more granular view of the age-association nominal p-values. Finally, the line plots show the combined motif match density across all distal or proximal peaks separated into significantly (FDR < 0.05) closing peaks in teal, opening peaks in brown, and not age-associated (nominal p-values > 0.5) peaks in grey. Each heatmap row represents an aggregate of approximately 150 OCRs.

### Enrichment of age-associated OCRs point to a remodeling of the skeletal muscle regulatory network

To test for enrichment of our age-associated OCRs in functional biological pathways, we linked our chromatin regions to genes via the nearest transcription start site (TSS) and then tested for enrichment of those genes in KEGG gene sets. Only four of the 213 tested KEGG terms were significantly (FDR < 0.05) enriched for age-associated peaks: pyruvate metabolism, RNA degradation, and propanoate metabolism in female type 1 fiber opening chromatin and Wnt signalling in male type 2x closing chromatin (Fig S6A) (Supplementary Table 4). While likely limited by the small number of age-associated OCRs and the difficulty of accurately associating OCRs with genes, these results may indicate that age-associated open chromatin regions are enriched for pathways involved in metabolism and cell-to-cell signalling.

To better understand the function of the age-associated OCRs, we tested for enrichment or depletion of age-associated OCRs in each chromatin state (Fig 3C; Fig 3D; Supplementary Table 5). This revealed a chromatin state-specific shift characterized by reduced accessibility to gene regulatory elements while keeping promoters and already transcribed genes accessible. Closing OCRs showed enrichment for active enhancers and depletion for active transcription start sites across sexes and fiber types (Fig 3D). Similarly, across sexes and fiber types, age-associated peaks trended towards depletion in transcribed states and enrichment in regions flanking transcription start sites (Fig S6B). These results suggest aging changes the accessibility of regulatory elements rather than directly suppressing transcribed regions or recruitment to promoters.

Finally, to identify transcription factors influencing or being impacted by age-associated epigenome remodeling, we tested for enrichment of more than 3,000 transcription factor motifs in age-associated OCRs. We identified 267 transcription factor motifs that were significantly (Bonferroni-adjusted p-value < 0.05) enriched across fiber types, sexes, and direction of association with age (Supplementary Table 6). NR3C1 (glucocorticoid receptor), NR3C2 (mineralocorticoid receptor), and NR3C4 (androgen receptor) were the most significantly enriched transcription factor motifs across all male fiber types, showing the highest nominal enrichment of any transcription factor in the closing chromatin of male type 2a fibers (Fig 3E). In female fiber types, nuclear factor I family members were the most significantly enriched transcription factor motifs, again showing the most significant enrichment in the closing chromatin regions of type 2a fibers (Fig 3E). We found that NR3C1 motifs were significantly (p-value < 1e-9) enriched for TSS distal (> 5kb) OCRs across all tested OCR sets from each fiber type / sex combination and that the motifs were significantly (p-value < 5.7e-17) more likely to occur in the center 150 base pairs of the OCR when compared to the flanking 150 base pairs (Fig 3F). We also observed a significant enrichment for CEBPB and ATF4 transcription factor motifs in the opening chromatin regions of both male and female type 2 fibers. Of the mentioned transcription factors, NFIA and CEBPB are significantly (FDR < 0.05) differentially expressed in our data, with NFIA showing a decrease in type 2a female fibers and CEBPB showing an increase in type 2a male fibers corresponding with their respective observed motif enrichment for closing and opening chromatin. Overall, these results suggest that age-associated changes in chromatin accessibility disproportionately overlap with NR3C1, NFIA, CEBPB, and ATF4 transcription factor motifs in a fiber-type and direction of age-association specific manner.

## Discussion

Here, we used the largest published human skeletal muscle biopsy single-nucleus RNA and ATAC dataset to date (N=287 donors) to explore sex-specific age-associated changes in *vastus lateralis* skeletal muscle nuclei composition, gene expression, and chromatin accessibility. Across 13 cell types, we identified 3 with age-associated differences in nuclei abundance. These findings, specifically the decrease in type 2x fiber and increase in neuronal and neuromuscular junction abundance, are in line with the findings of prior literature which characterize an age-related increase in denervation prompting reinnervation attempts and preferential type 2 fiber loss^4,15,16,25–28^. We did not observe previously reported changes, including decreased endothelial, satellite, and smooth muscle cell nuclei^13,29^ or increased type 1 fiber^15^ and FAP^29^ nuclei. These differences in findings are likely due to both a difference in sample size (287 vs <23 single-cell samples) and a difference in the age range of the respective cohorts. The participants of our study are predominantly between the ages of 50 and 80 and while some of the other studies compare individuals in their 20s to those in their 70s^13,29^.

We identified hundreds of age-associated genes and thousands of age-associated OCRs, which were enriched in a small number of KEGG pathways. When comparing DEGs found in our study to those identified in a prior study by Kedlian et al.^16^ we found low (58-64%) but significant (highest nominal p-value = 0.01 among all three fiber types by two sided Fisher’s Exact Test) agreement in the direction of aging association effect when compared to a null assumption of 50% agreement. However, possibly due to both the difference in sample size (287 this study vs 17 for Kedlian et al.^16^ single cell samples) and biopsy site (vastus lateralis for this study vs intercostal muscle for Kedlian et al.^16^), saw very little (16%) overlap of significant DEGs in our study. Age-associated molecular features in one sex or fiber type generally showed the same direction of effect across sex or fiber types (even if not reaching statistical significance), suggesting similarity in aging as previously seen in a bulk RNA-seq study performed by de Jong et al^14^. This similarity in age-associated changes to molecular features was also reflected in a concordance of KEGG term enrichment across fiber type and sex. For example, the cellular senescence KEGG term leaned towards enrichment in all sexes and fiber types consistent with findings in previous studies^13–16^. ABC transporters and glycerolipid metabolism also showed consistent direction of effect, but have not appeared in past enrichment analyses. In contrast, our most significantly enriched term, proteasome, showed age-associated gene enrichment in female type 1 fiber while trending towards depletion in male fiber types. This enrichment was driven by significant increases in the expression of two proteasome subunits and a nominal increase in the expression of eight other subunits. While excessive protein catabolism can result in muscle wasting, prior literature shows decreased proteasome activity associated with both older and type 2 muscle fibers, and low responsiveness of the ubiquitin-proteasome system may result in impaired clearance of damaged or misfolded proteins^15,25,27,30,31^.

While most KEGG terms significantly enriched for age-associated genes and OCRs leaned towards enrichment (even if non-significant) across sex and fiber type within a modality, there was no overlap in significantly enriched or depleted terms between the modalities. This may be due to the small number of terms enriched for age-associated OCRs or technical difficulties linking regions of open chromatin to the genes they regulate. It may also reflect a difference in information encoded by modalities; for example, studies have shown that the epigenome may prime a cell to respond to environmental changes without inducing immediate changes in transcription^19,32,33^.

Prior studies using ATAC and methylation data from a variety of human tissues have yielded conflicting reports on whether increased accessibility, decreased accessibility, or neither characterises the aging process^34–39^. Overall, we identified more closing chromatin regions than opening ones. We note that our data may not allow us to accurately assess extremely widespread, global changes in chromatin structure with aging, as such changes may be hidden by standard library size normalization techniques.

We found closing chromatin was enriched in enhancer regions but depleted in active TSS regions. Age-associated chromatin, both opening and closing, was also enriched in TSS flanking regions. Though this is a novel finding for human skeletal muscle, prior studies of age-associated OCRs in human blood and liver samples reported enrichment in enhancer and promoter chromatin states, but gave conflicting results for enrichment in closing versus opening chromatin^35,37–39^. Our results suggest that aging may impact skeletal muscle through altered accessibility of regulatory elements outside of transcribed regions and their respective promoters. Past GWAS have found that genetic variants associated with skeletal muscle and age-related diseases such as T2D reside predominantly outside of protein-coding regions; this finding supports a potential mechanism by which these variants may interact with aging pathology^40^.

Testing for TF motif enrichment within age-associated OCRs yielded significant enrichments across sex, fiber type, and direction of association with age, with the most significant result being the enrichment of NR3C1 motifs and those of its family members, in male type 2 closing chromatin. This suggests that reduced signalling from these nuclear receptors, including the glucocorticoid, mineralocorticoid, and androgen receptors, may contribute to or result from the aging process. While chronic exposure to glucocorticoids can drive muscle wasting^41^, prior literature also shows NR3C1 expression is positively correlated with muscle mass in non-sarcopenic patients^42–44^. While we observed no age-associated change in NR3C1 expression, NR3C1’s role in acute responses to tissue damage may not be evident in the RNA of baseline resting muscle biopsies. The decrease in accessibility of NR3C1 motif regions may, however, indicate a cell state of reduced glucocorticoid signalling sensitivity. Other regulatory mechanisms, such as low-functioning NR3C1 isoforms and cortisol binding competition, may influence NR3C1 activity without altering transcript levels^42,44,45^. Although glucocorticoid receptor binding impacts multiple pathways, evidence suggests that it can promote ubiquitin–proteasome formation and the healthy catabolism necessary for muscle tissue regeneration and repair^41,42^. The other significantly enriched family members, NR3C2 and NR3C4, also regulate skeletal muscle maintenance through hormone binding and the promotion of muscle regeneration^46–48^. It is worth noting that both NR3C1 and NR3C4 show sex-specific activity, especially the androgen receptor NR3C4, which may contribute to the male specificity of NR3C1/4 motif enrichment^17,47,49^. We also identified TF motifs enriched in closing chromatin in females, NFIX and its family members, and in opening chromatin across fiber types and sexes, ATF4 and CEBPB. Though these changes in motif accessibility are not always reflected in a change in transcription factor expression, prior literature evidences a role for NFIX, ATF4, and CEBPB in managing human skeletal muscle atrophy^12,50–53^. These motif enrichments were often highly significant and demonstrate that changes in the epigenome may not be apparent in matched transcriptomes, showing the importance of chromatin accessibility analyses.

Our study has several limitations. First, while donors ranged between 20 and 79 years of age, 80% aged between 50 and 70. This limits our ability to detect changes outside this range and makes comparison to prior literature inexact, as these often compare individuals under 30 to individuals over 50^13,15,16^.

Second, we focused on muscle fiber types for most analyses, as statistical power to detect age-associated molecular features in cell types with fewer nuclei was limited. Third, this study, like most previous studies, is observational, making potential adjustment for generational differences in lifestyle difficult when comparing older and younger individuals. Fourth, biopsy location also complicated cross-study comparison. While this and many studies use *vastus lateralis* biopsies, others use arm or intercostal muscle biopsies, which may experience different lifetime exercise exposure, which has a strong effect on muscle aging^18^.

In summary, we replicated known age-associated changes in skeletal muscle composition, and identified novel biology in the overlap between age-associated closing chromatin regions and regulatory elements in skeletal muscle fibers. In particular, we found that age-associated closing chromatin regions were enriched for enhancers while being depleted for active transcription start sites. Further, we found evidence for a mechanism whereby age-associated closure of OCRs with hormone-binding TF motifs may reduce sensitivity to signals regulating muscle maintenance and regeneration, leading to sex- and fiber type-specific atrophy.

## Methods

### Sample collection and data generation

Sample collection in the Finland-United States Investigation of NIDDM Genetics (FUSION) Tissue Biopsy Study is described in Scott et al^54^. The methods for performing snRNA-seq and snATAC-seq on the FUSION samples is described in Varshney et al^19^. Briefly, 287 frozen muscle tissue biopsy samples were organized into 10 batches balanced by sex, cohort, age, BMI, and oral glucose tolerance test (OGTT) result. The resulting sequencing libraries were mapped to the hg38 human genome reference and subject to a QC pipeline to identify doublets and remove contamination before jointly clustering the RNA and ATAC nuclei using Liger (1.0.0)^55^. These clusters were then annotated with canonical marker gene expression for each of the cell-types and cell type- and participant-specific bam files were created^19^.

### ATAC-seq peak calling and consensus peak feature definition

We called peaks, defined consensus peaks, and defined nuclei type-specific OCRs as described in Varshney at al^19^. Briefly, we used MACS2 (2.1.1.20160309)^56^ to call peaks and filtered those peaks according to a stringent significance threshold (0.1% FDR). We then defined a consensus peak set across all 13 clusters by collapsing summits within 150bp and retaining those with the greatest significance resulting in 983,155 consensus summits of 301bp in length. We used the tau metric from Tanai et al.^57^ to define cell type-specific OCRs for use in downstream analysis.

### Differential nuclei type abundance

Using 285 FUSION samples (121 female, 164 male) with nuclei of all 13 types we used DESeq2 (1.46.0)^20^ to test for association between total combined ATAC and RNA nuclei counts and age for each cell type independently while adjusting for batch, BMI, OGTT, and applying an offset for size factors. Only associations with FDR < 5% were counted as significant to adjust for multiple testing across cell types and sexes.

### Single-nucleus differential gene expression with respect to age (RNA)

For all cell types, except adipocytes and T-cells which were excluded for having fewer than 500 nuclei, and in combined, male, and female sample sets, we tested genes with at least one count in at least 10% of nuclei for differential expression with respect to age. Using the MAST (1.32.0)^58^ R (4.2) package, we applied a linear mixed model with a random effect term for the donor to log adjusted TPM gene count data while adjusting for batch, BMI, OGTT, cellular detection rate^58^ and sex in the combined sex analysis. We used the ‘glmer’ method from the lme4 (1.1_35.5)^59^ R package and set the ‘ebayes’ and ‘strictConvergance’ options to false and nAGQ to 0. For comparison, we also tested the association between pseudobulked gene expression and age using DESeq2 with adjustments for BMI, number of nuclei, batch, OGTT, and an offset term for library size; for the DESeq2 analysis we tested genes with at least 10 counts in 10% of samples. To account for multiple testing, we used a threshold of FDR < 5% across all tested genes within each cell type and sex combination when using either the single-nucleus or pseudobulk method.

### Comparison of differential expression results to Kedlian et al

LTSR > 0.9 genes for type 1 and 2 fibers and fragments were taken from the supplementary tables available in Kedlian et al^16^. Differentially expressed genes were combined to make type 1 and type 2 lists which were then compared to the closest corresponding list from our own data (type 1 with type 1 and type 2 with both type 2a and type 2x). As nominal p-values were not available for the significant genes from Kedlian et al we only quantified the overlap in significance (FDR < 0.1) in our data and the directional concordance of coefficients. Significance of agreement in direction between significantly age-associated genes from Kedlian et al. and results from this study was performed via two sided Fisher’s exact test with the null assumption being 50% agreement in direction of effect.

### Gene set enrichment analysis for muscle fiber transcriptome

For each fiber type and sex we tested KEGG terms for enrichment with age-associated genes using the method described in the RNA-Enrich paper^21^. Briefly, this method tests each term for association between the gene membership within a term and the nominal p-value output from the differential expression analysis. We tested only KEGG terms containing between 10 and 1,000 genes, ignoring overly specific terms which would show strong results even if only a single gene was age-associated and overly general terms which would be less informative to see enrichment in. To account for multiple testing, we used a threshold of FDR less than 5% across all tested KEGG terms within each fiber type and sex combination.

### Single-nucleus differential chromatin accessibility with respect to age (ATAC)

For each cell type, and in combined, male, and female samples, we tested our previously identified cell type-specific OCRs for differential accessibility with respect to age. We tested only OCRs having at least 1 count in at least 10% of nuclei. Using a generalized linear mixed model with a negative binomial distribution^60^ we tested for association between OCR counts and age while applying a random effect term for donor, an offset term for total counts and adjusting for batch, BMI, OGTT, TSS enrichment, and sex for the combined analysis. We used the DHARMa (0.4.7)^61^ package tools for reporting changes to Akaike’s information criterion and zero-inflation and inform our decisions about covariate inclusion and distribution for our generalized linear mixed model respectively. We also referenced Teo et al for best practices when testing for differential accessibility at the single-nucleus level.^62^ For a more direct comparison on our data, we also tested the association between pseudobulked OCR counts and age using DESeq2 with adjustments for batch, bmi, log10 number of nuclei per sample, OGTT, TSS enrichment, and sex for the combined analysis. For the pseudobulk analysis, we tested OCRs with at least 10 counts in 10% of samples. To account for multiple testing, we used a threshold of FDR < 5% across all tested genes within each cell type and sex combination when using either the single-nucleus or pseudobulk method.

### Gene set enrichment analysis for muscle fiber epigenome

For each fiber type, sex, and chromatin behavioral pattern, we identified KEGG terms enriched for age-associated OCRs using the chipenrich^63^ R package. We used the nearest TSS method to link our list of age-associated peaks to genes after which chipenrich (2.30.0) tested for enrichment of those linked genes with member genes of KEGG terms. We tested only KEGG terms containing between 10 and 1,000 genes, ignoring overly specific terms which would show strong results even if only a single gene was age-associated and overly general terms which would be less informative to see enrichment in. We then removed any terms that showed up as enriched in a background of all peaks tested for age association within a given cell type and sex combination. To account for multiple testing, we used a threshold of FDR < 5% across all tested KEGG terms within each fiber type and sex combination. Chromatin state enrichment analysis for muscle fiber differentially accessible regions:

For each fiber type, sex, and age association direction of effect (opening or closing) we used Roadmap Epigenomics^64,65^ chromatin states for adult human female skeletal muscle (E108) and adult human male skeletal muscle (E107) to identify enrichment of age-associated OCRs in specific chromatin states. We assigned each OCR to a chromatin state based on the center of the OCR, and then tested the chromatin states “Active_TSS”, “Flanking_TSS”, “Flanking_transcription”, “Strong_transcription”, “Weak_transcription”, “Genic_enhancer”, “Active_enhancer”, “ZNF_gene_repeats”, “Heterochromatin”, “Bivalent_TSS”, “Flanking_bivalent”, “Bivalent_enhancer”, “Repressed_polycomb”, “Weak_repressed_polycomb”, and “Quiescent_low_signal” for enrichment of age-associated OCRs. We used a logistic regression model with a background of OCRs with a nominal p-value of 0.5 or greater for age-association in the tested fiber type to ensure they were truly not age-associated, and adjusted for OCR GC content, median counts per nucleus, and a binary term for being distal (within 5kb) from the nearest TSS. To account for multiple testing, we used a threshold of Bonferroni-adjusted p-value < 0.05 across all chromatin states, sexes, fiber types, and age-association direction of effect.

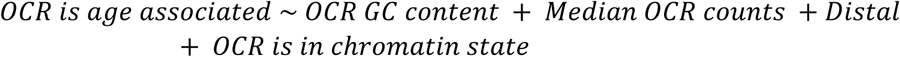

Upon comparison, the results from adult female and adult male skeletal muscle Roadmap chromatin states appeared very similar, so we report the female results in figures 3C and 3D, but the complete results using both Roadmap references can be found in supplementary figure 6B along with results from two female fetal muscle references.

### Motif enrichment analysis

For each fiber type, sex, and age association direction of effect (opening or closing) we used a previously curated set of 3110 transcription factor motifs^66^. We assigned OCRs to transcription factor motifs based on the center of the OCR, and then tested each transcription factor motif for enrichment of age-associated OCRs. We used a logistic regression model with a background of OCRs with a nominal p-value of 0.5 or greater for age-association in the tested fiber type, and adjusted for OCR GC content, median counts per nucleus, and a binary term for being distal (within 5kb) from the nearest TSS. To account for multiple testing, we used a threshold of Bonferroni-adjusted p-value < 5% across all tested transcription factor motifs, sexes, fiber types, and age association direction of effect.

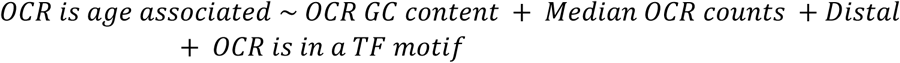

Testing the relative frequency of peaks with a motif match in the central 150 base pairs compared to the 150 base pairs flanking that central region was done via Fisher’s exact test.

Supplemental 1:

a.) Per sex and fiber type volcano plots of differential gene expression with respect to age. Genes with FDR < 0.05 for age-association have been colored red.

Supplemental 2:

a.) Signed -log10 p-values (+ higher expression with age, - lower expression with age) of differential expression in each sex between each fiber type analysis in a pairwise arrangement. The number and percentage of plotted genes are recorded in the corners of each scatterplot.

b.) Signed -log10 p-values (+ higher expression with age, - lower expression with age) of differential expression in each fiber type and sex between the single-nucleus level MAST analysis and the pseudobulk DESeq2 analysis. The number and percentage of plotted genes are recorded in the corners of each scatterplot.

Supplemental 3:

a.) Signed -log10 p-values from this study compared to signed -log10 1-ltsr (local true sign rate) values from Kedlian et al.^16^.Infinite -log10 1-lstr values resulting from a rounded up lstr value of 1 reported in Kedlian et al. are plotted as if a value of 0.9999999999 was reported instead for the purposes of plotting. The number and percentage of plotted genes are recorded in the corners of each scatterplot.

Supplemental 4:

a.) Per sex and fiber type volcano plots of differential open chromatin region accessibility with respect to age. OCRs with FDR < 0.05 for age-association have been colored red.

Supplemental 5:

a.) Signed -log10 p-values (+ more accessibility with age, - less accessibility with age) of differential chromatin accessibility in each sex between each fiber type analysis in a pairwise arrangement. The number and percentage of plotted OCRs are recorded in the corners of each scatterplot.

b.) Signed -log10 p-values (+ more accessibility with age, - less accessibility with age) of differential chromatin accessibility in each fiber type and sex between the single-nucleus-level generalized linear mixed model analysis and the pseudobulk DESeq2 analysis. The number and percentage of plotted OCRs are recorded in the corners of each scatterplot.

Supplemental 6:

a.) Heatmap of KEGG terms with FDR < 0.05 enrichment or depletion in at least one set (Sex, fiber type, and direction of age-association combination) of age-associated OCRs. Squares for enrichment tests in which zero overlaps between age-associated OCR-linked genes and gene set genes are detected have been grayed out.

b.) Heatmap of 15 chromatin states from the Roadmap database showing enrichment or depletion of age-associated OCRs in 4 Roadmap database samples representing adult female and male skeletal muscle, and two from fetal female muscle.

