## Supplementary figures and images for "Erosion of regenerative regulation: age-associated shifts in the skeletal muscle fiber epigenome and transcriptome"

### Supplemental Figure 1

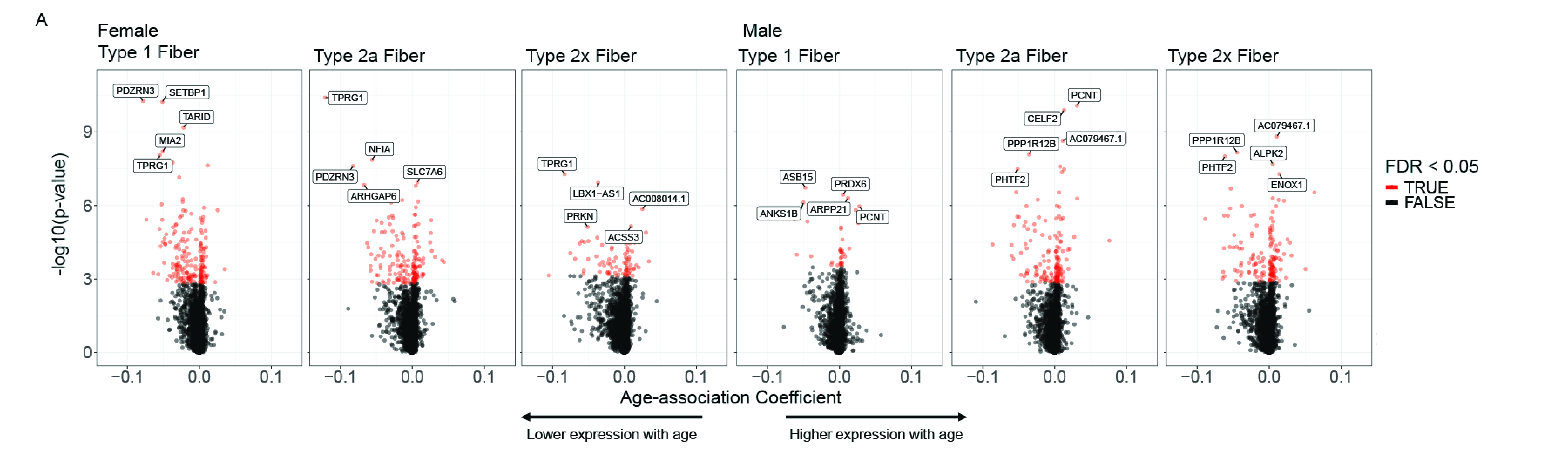

### Supplemental Figure 2

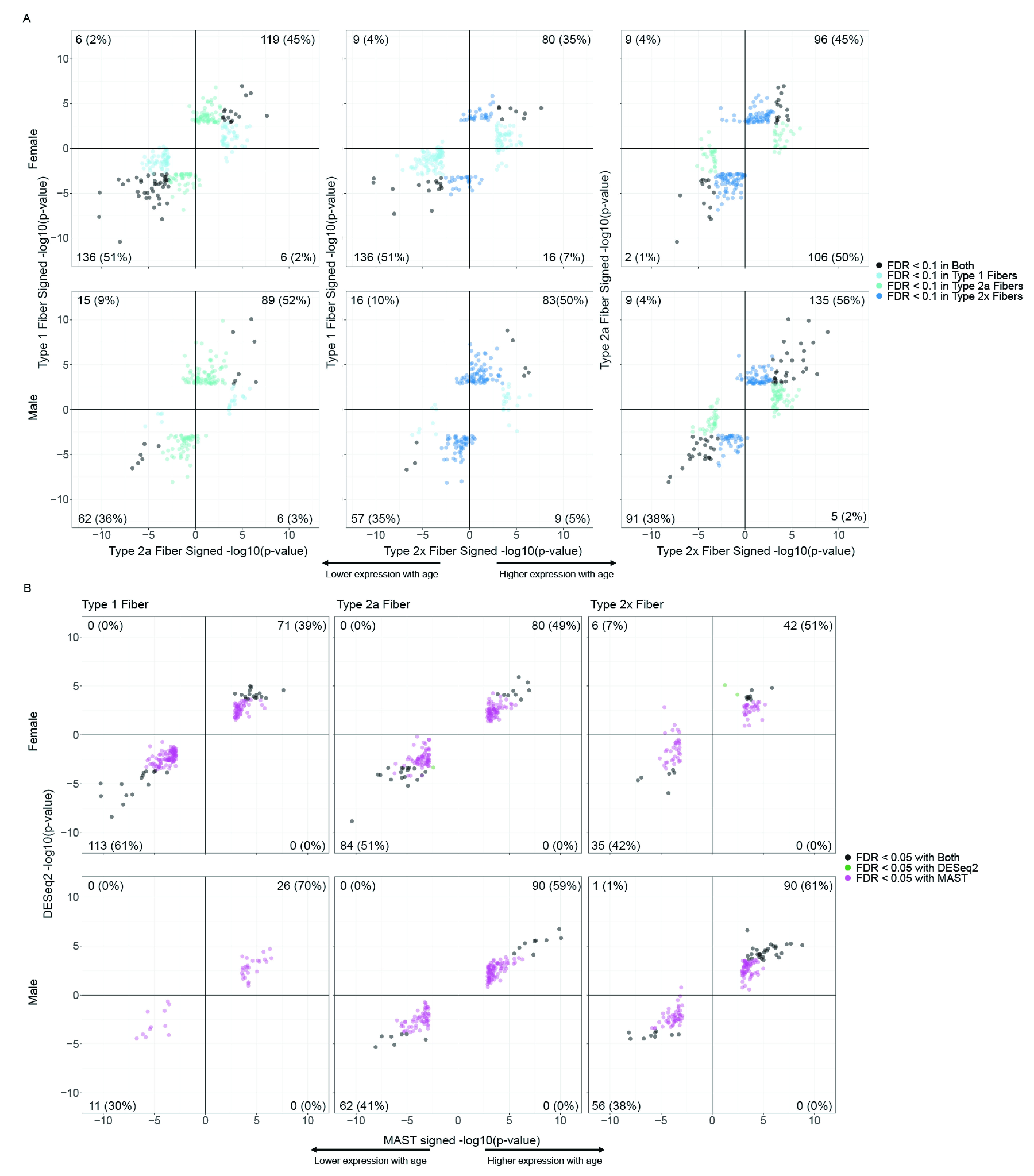

### Supplemental Figure 3

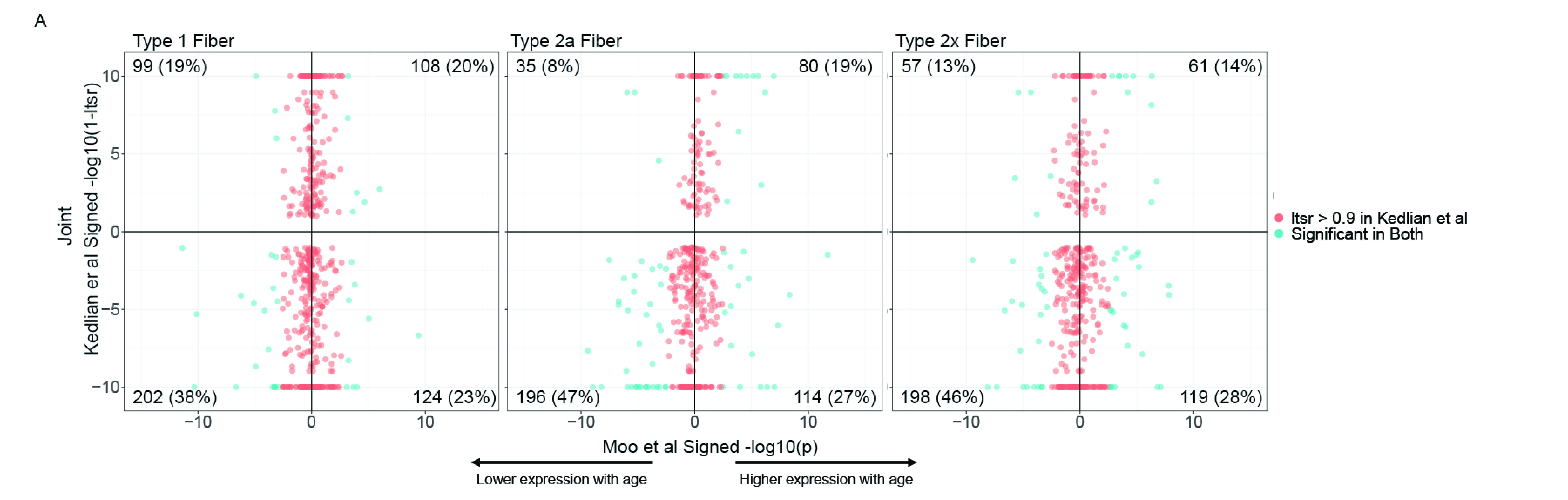

### Supplemental Figure 4

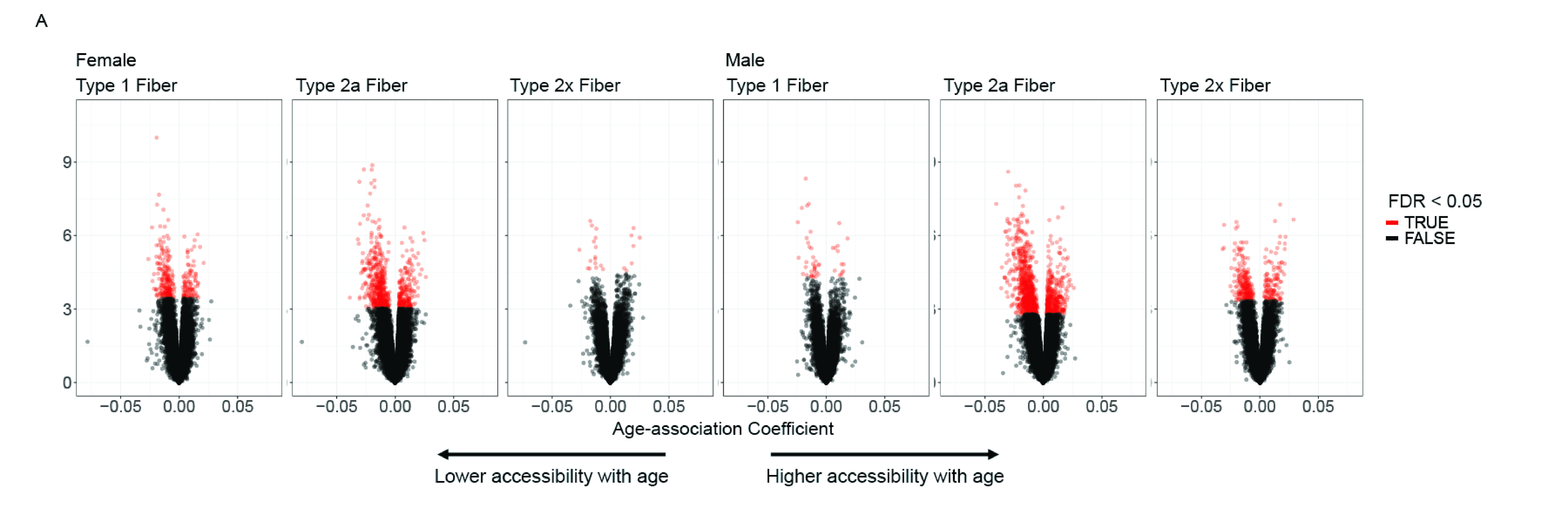

### Supplemental Figure 5

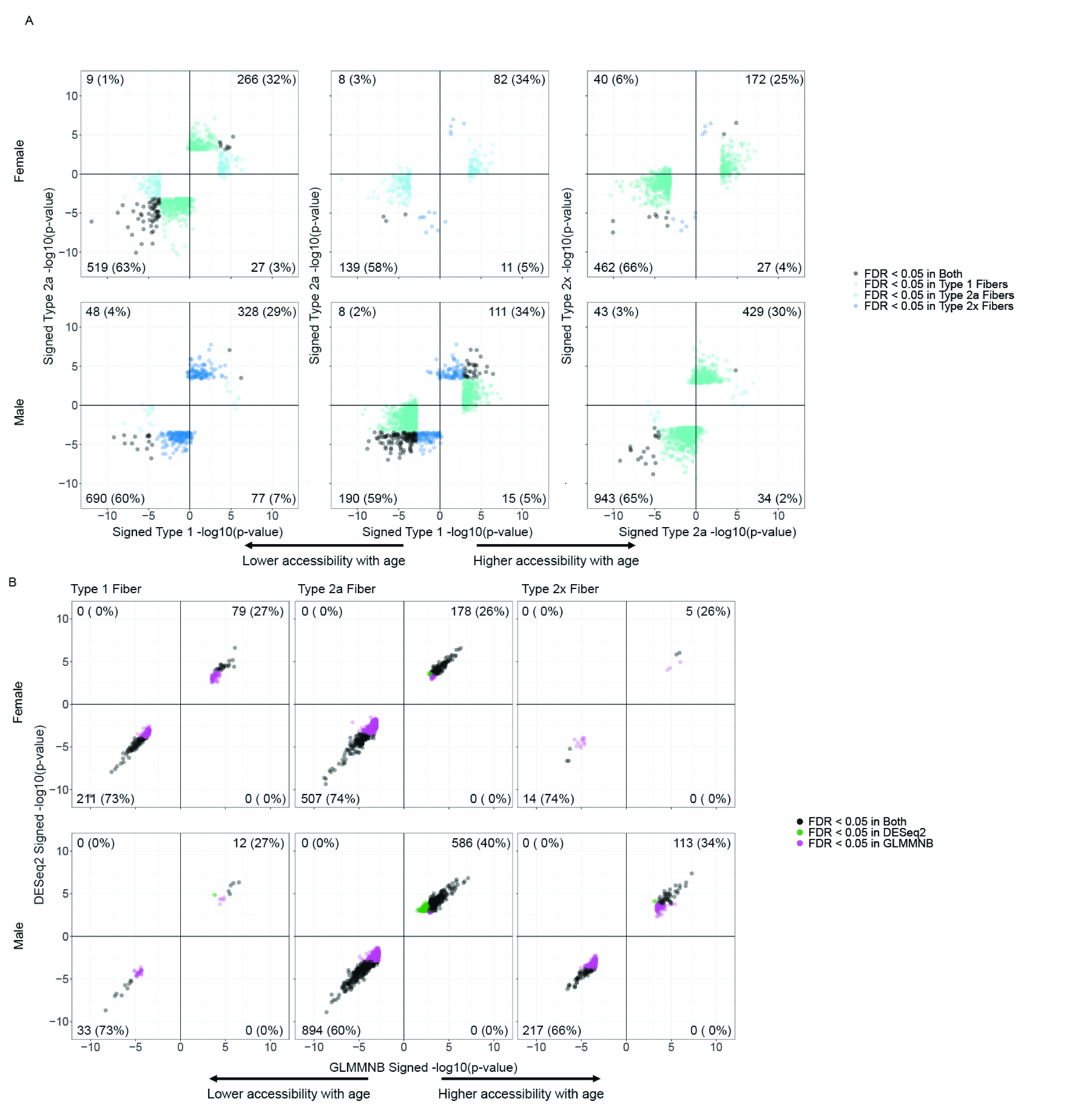

### Supplemental Figure 6

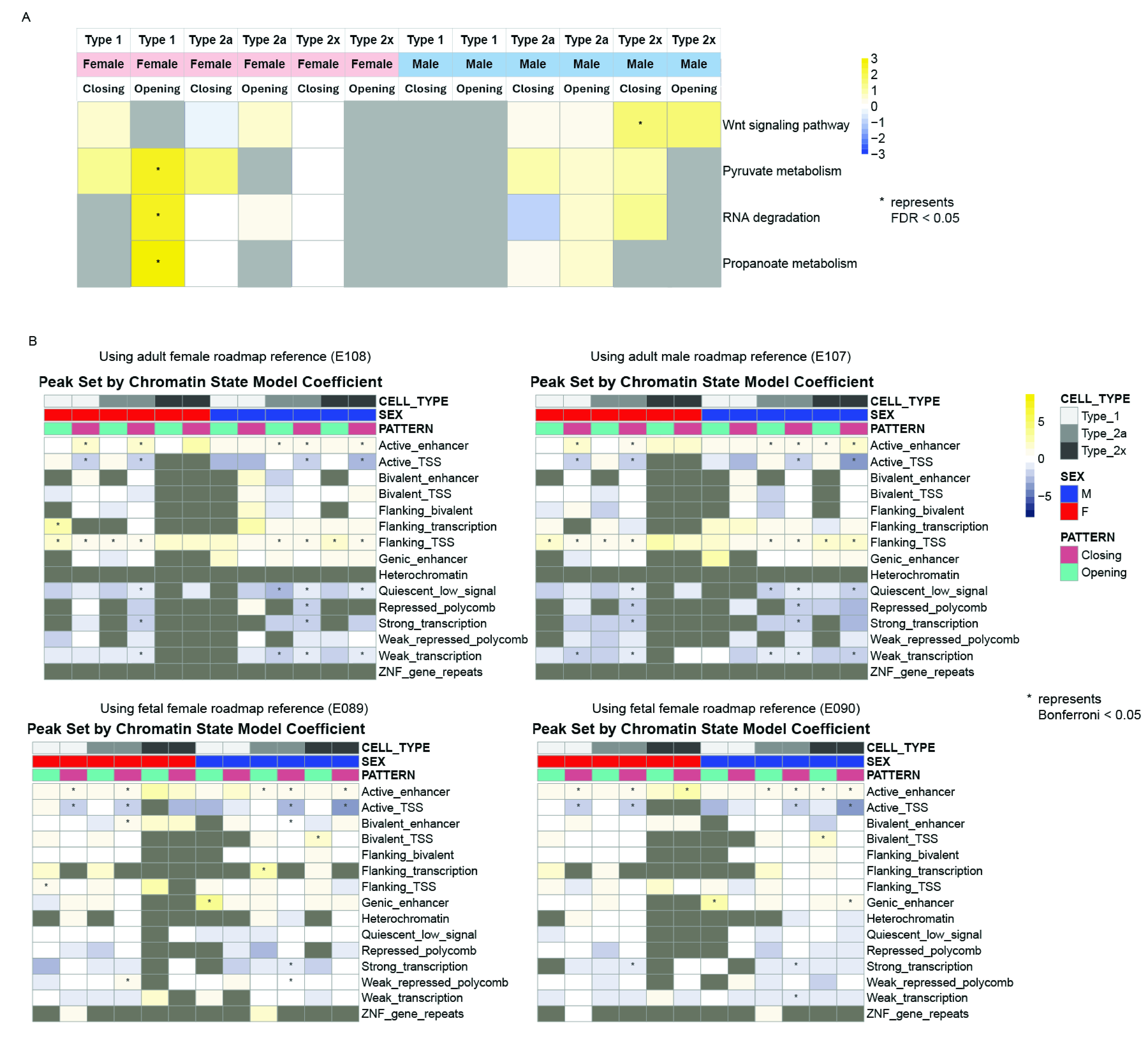
